# Imaging Three-Dimensional Brain Organoid Architecture from Meso- to Nanoscale across Development

**DOI:** 10.1101/2021.12.03.471084

**Authors:** Juan Eduardo Rodriguez-Gatica, Vira Iefremova, Liubov Sokhranyaeva, Si Wah Christina Au Yeung, Yannik Breitkreuz, Oliver Brüstle, Martin Karl Schwarz, Ulrich Kubitscheck

**Affiliations:** Institute of Physical and Theoretical Chemistry, University of Bonn, Wegelerstr. 12, 53115 Bonn, Germany; Institute of Reconstructive Neurobiology, University of Bonn, Sigmund-Freud-Strasse 25, 53127 Bonn, Germany; Institute Experimental Epileptology and Cognition Research (EECR), University of Bonn Medical School, Sigmund-Freud-Str. 25, 53127 Bonn, Germany; LIFE & BRAIN GmbH, Cellomics Unit, Venusberg-Campus 1, D-53127 Bonn, Germany

**Keywords:** brain organoid, expansion microscopy, light sheet fluorescence microscopy, super-resolution, synaptogenesis

## Abstract

Organoids are human stem cell-derived three-dimensional cultures offering a new avenue to model human development and disease. Brain organoids allow studying various aspects of human brain development in the finest details *in vitro* in a tissue-like context. However, spatial relationships of subcellular structures such as synaptic contacts between distant neurons are hardly accessible by conventional light microscopy. This limitation can be overcome by systems that quickly image the entire organoid in three dimensions and in super-resolution. To that end we have developed a setup combining tissue expansion and light sheet fluorescence microscopy for imaging and quantifying diverse spatial parameters during organoid development. This technique enables zooming from a mesoscopic perspective into super-resolution within a single imaging session, thus revealing cellular and subcellular structural details in three spatial dimensions, including unequivocal delineation of mitotic cleavage planes as well as the alignment of pre- and postsynaptic proteins. We expect light sheet fluorescence expansion microscopy (LSFEM) to facilitate qualitative and quantitative assessment of organoids in developmental and disease-related studies.

**Summary statement:** The combination of light sheet fluorescence and expansion microscopy enables imaging of mature human brain organoids *in toto* and down to synaptic resolution

## Introduction

In the past years, advances in stem cell technologies enabled rapid progress in the field of pluripotent stem cell-based three-dimensional (3D) cultures such as brain organoids or spheroids. These culture formats represent self-organizing structures that recapitulate certain aspects of in vivo brain development. They display complex structures that recapitulate several aspects of early neurogenesis, including the formation of an apical and basal surface, polarized neuroepithelium, neurogenic ventricular and outer radial glia (oRG), the formation of layered, cortex-like architectures and maturation to the level of synapse formation (Paşca et al., 2015; Lancaster et al., 2017). At the same time, proper visualization of these diverse processes in 3D has remained challenging, and most analyses of their cyto- and histoarchitecture are still based on conventional two-dimensional (2D) histology of organoid cryosections. However, recently more sophisticated approaches for whole-organoid clearing and imaging that allow addressing the structural features in all three dimensions and preserving the 3D organization by avoiding cryosectioning were introduced and showed rapid development (Adhya et al., 2021; Albanese et al., 2020; Edwards et al., 2020). Whereas several studies that employ whole-organoid clearing have been published in the last years, they mainly use small organoids, e.g. to assess early neural differentiation (Benito-Kwiecinski et. al. 2021). A number of parameters remain rather challenging in the context of whole-mount analysis of larger organoids. This applies particularly to structures appearing at late stages of organoid differentiation such as imaging of dendritic spines and synapses (Masselink et al., 2019; Dekkers et al., 2019; Albanese et al., 2020).

Parallel to the development of organoids, novel, and fast large volume imaging methods were introduced that are able to depict fine cellular and subcellular structural features within geometrically extended tissue samples in 3D. The task to image large neuronal tissue fragments came into reach with the advent of Light Sheet Fluorescence Microscopy (LSFM). This technique allows to observe a fluorescently labeled specimen with one or two microscope objective lenses, whose focal plane is illuminated perpendicular to the detection axis by a thin sheet of light (Huisken et al., 2004; Dodt et al., 2007). Thus, LSFM offers intrinsic optical sectioning, which can further be amended by confocal line detection (Silvestri et al., 2012; Baumgart and Kubitscheck, 2012) to yield optimal contrast in scattering specimen. Substantial progress has been made in the past few years by using illumination beam shaping to achieve very thin light sheets to enhance optical resolution over large fields of view. Lattice light sheets (Chen et al., 2014; Ellefsen and Parker, 2018; Stockhausen et al., 2020), Airy beams (Vettenburg et al., 2014), or modifying the Gaussian light-sheet waist across the field of view (Dean et al., 2015; Fu et al., 2016; Neyra et al., 2020) resulted in extra thin light sheets with large extensions. 3D image stacks can be acquired by moving the specimen through the illuminating sheet within the detection plane of the imaging objective. When using sensitive and fast CMOS cameras, image rates of hundreds of frames per second can be achieved, significantly decreasing the imaging time for large specimens compared to confocal laser scanning microscopy.

Optimal usage of light-sheet microscopy requires the elimination of refractive index inhomogeneities in the probe by using an immersion medium with a refractive index matched to the cellular components of the probe. This can be accomplished by tissue *clearing* (for review on different clearing procedures, see Ueda et al., 2020). Combining tissue clearing with LSFM allows effective optical resolutions in the range of 0.3 μm laterally and 1.0 μm axially when using long-distance objective lenses for imaging with a numerical aperture (NA) of 1.0 or greater. Thus, LSFM is especially well suited for the fast analysis of complex arrangements of large cleared cell clusters and tissue fragments, where it enables, e.g., fast light microscopic assessment of the complex 3D architecture of organoids and spheroids (Albanese et al., 2020; Benito-Kwiecinski et al., 2021).

However, LSFM cannot reveal the very fine details of neuronal networks since these structures are well below the optical diffraction limit. To visualize synapses with spatial information conserved in a 3D human cerebral organoid, super-resolution imaging is necessary. Classical point scanning light microscopy in super-resolution is challenging and restricted to small regions (single synapses), making it virtually impossible to scan entire organoids or even distinct structures within organoids in 3D.

A solution to this issue came into reach with the development of light-sheet fluorescence expansion microscopy (LSFEM), which has enabled the analysis of extended neural circuits in super-resolution (Bürgers et al., 2019; Gao et al., 2019; Schwarz and Kubitscheck, 2021). As in standard Expansion Microscopy (ExM, Chen et al., 2015; Ku et al., 2016; Tillberg et al., 2016; Chozinski at al., 2016), LSFEM utilizes water-absorbent polymers to physically expand enzymatically treated tissue samples. Before synthesizing the expandable polymer within the fixed tissue sample, proteins of interest are labeled with fluorescent antibodies that bind the antigen of interest. The tissue is then partially digested to allow subsequent expansion of the polymer matrix containing the fluorescent labels. As a result of expansion, fluorescent moieties spaced closer than the optical diffraction limit (about 250 nm) can be optically resolved, resulting in effective super-resolution images of organoids. Using LSFEM, we were able to rapidly image extended neuronal circuits in effective super-resolution (Bürgers et al., 2019).

Here we present a novel brain organoid analysis pipeline, which employs LSFM and LSFEM to image entire brain organoids during different developmental stages in 3D. The methods allows to zoom in and out an entire single organoid from meso-to nanoscale optical resolution in order to obtain a comprehensive view of the both the brain organoid architecture and subcellular aspects. Careful sample preparation allowed the conservation of fluorescent proteins, which are frequently used to label e.g. neuronal subpopulations. Using effective super-resolution imaging, the finest details of neuronal network parameters within a larger context can be depicted. Ultimately, we succeeded for the first time to identify clusters of synaptically connected neurons in the context of an entire cleared mature brain organoid.

## Results

### Clearing, physical expansion and LFSM enable the analysis of mature brain organoids

Fixed brain organoids represent opaque structures. We developed specific sample preparation techniques and imaging approaches to exploit the inherent information optimally (see Methods). Following fixation and permeabilization, we usually employ DNA staining to mark all cell nuclei. We also include markers for specific cell types or cellular structures highlighting specific types of neurons or subcellular neuronal structures. To allow a light microscopic analysis of these samples, optical clearing is mandatory (Fig. 1A). Therefore, the first steps of ExM include the addition of bifunctional linker coupling protein residues, thereby creating a polyacrylamide gel within the organoid, which keeps the fluorescently labeled structures in place. Digestion by proteinase K renders the sample transparent. Keeping the sample in a buffered aqueous immersion medium such as PBS induces a 1.5-fold expansion (Fig. 1B), whereas the exchange of the medium to bidistilled water results in an approximately 4-fold physical expansion (Fig. 1C). In either state, the sample can be analyzed using LSFM (Fig. 1D).

**Figure 1.**
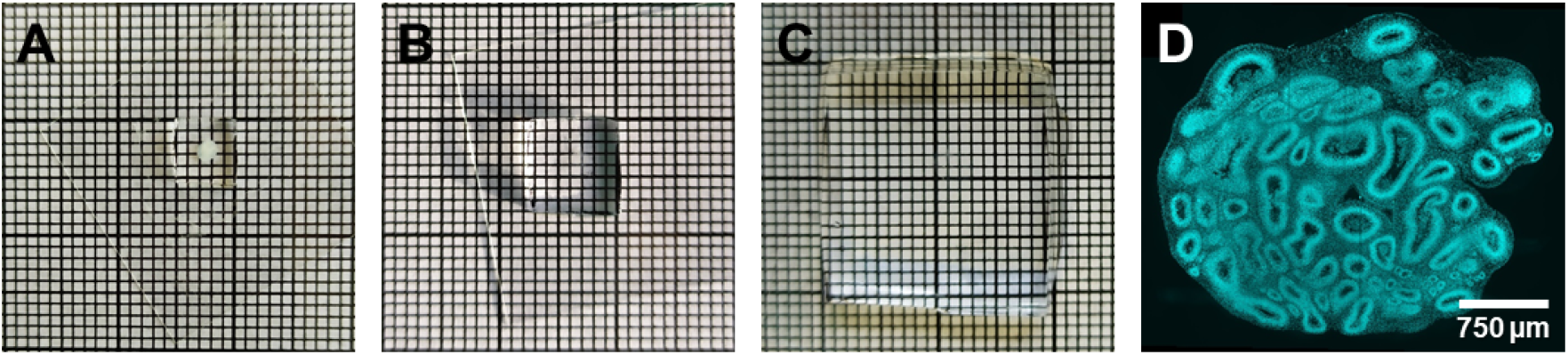
Brain organoid before and after clearing and expansion. (A) Two-month-old brain organoid embedded in a polyachrylamide gel. (B) Two-month-old organoid after proteinase K digestion, which resulted in a clearing of the organoid and an approximately 1.5-fold expansion. (C) The same organoid after expansion in bidistilled water, which yielded an approximately 4-fold expansion. (D) Optical section of the cleared and 1.5-fold expanded organoid showing the cell nuclei of an optical section in a depth of 1.2 mm.

#### LSFM enables analysis of brain organoid structures across development and cell differentiation

##### Tracing subpopulations of fluorophore-labeled cells

Mesoscale imaging of cleared brain organoids allowed to follow organoid development starting from the time point of generation until maturation. The key steps of the pipeline for organoid sample preparation and imaging were described in Fig. 2, and the various approaches to imaging summarized in Table 1. Cerebral organoids can reach extensions of up to several millimeters. To cover such dimensions, light-sheet imaging was performed using a low magnification objective lens (10x) with a relatively low numerical aperture (NA, 0.3) in order to achieve a large field of view for covering the complete organoid with a limited number of mosaic tiles.

**Figure 2.**
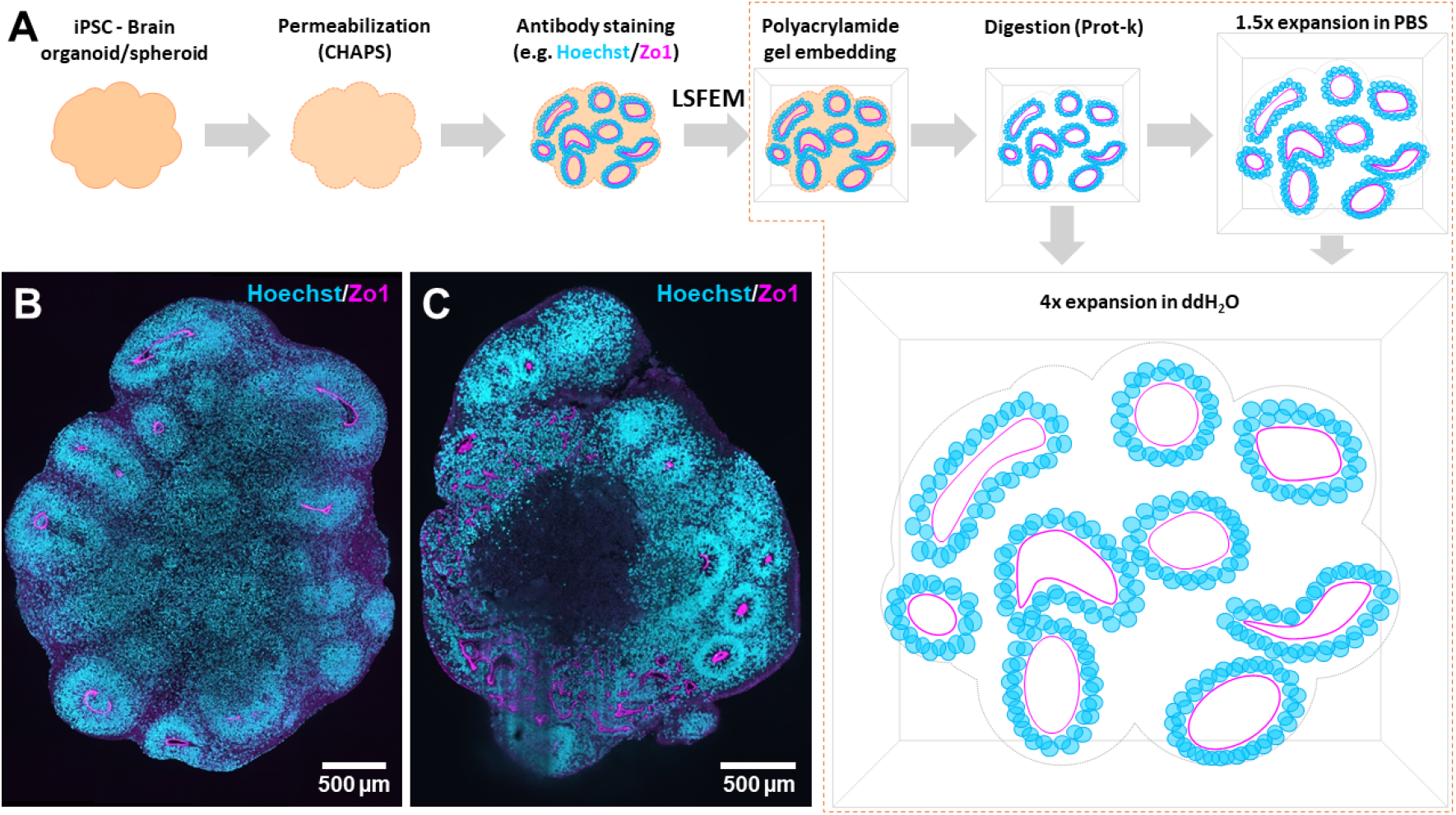
Organoid sample preparation for LSFEM. (A) Pipeline for organoid sample preparation. After fixation of the 3D samples, a permeabilization step is made using a detergent (CHAPS), which is a key step to allow proper whole-body immunostaining of the organoid. Subsequently, immunostaining for identifying specific cell types or structures, along with a nuclear staining is performed. In the diagram, the tight junction marker Zo1 – magenta, and nuclear staining Hoechst – cyan, are used as example. After immunostaining the sample is embedded in and chemically linked to a polyacrylamide gel. A digestion or mechanical homogenization of the sample using a buffer containing proteinase K (Prot-K) renders the sample transparent, resulting in optimal conditions for light-sheet imaging. Placing the sample after digestion in PBS leads to an isotropic expansion of 1.5-fold, while placing the digested sample in bi-distilled water leads to a 4-fold expansion, allowing for the analysis of the whole organoids or spheroids in super-resolution. Two optical sections of (B) a three-month-old brain spheroid and (C) a two-month-old brain organoid are shown as an example. Both are stained against Hoechst and Zo1.

**Table 1.**
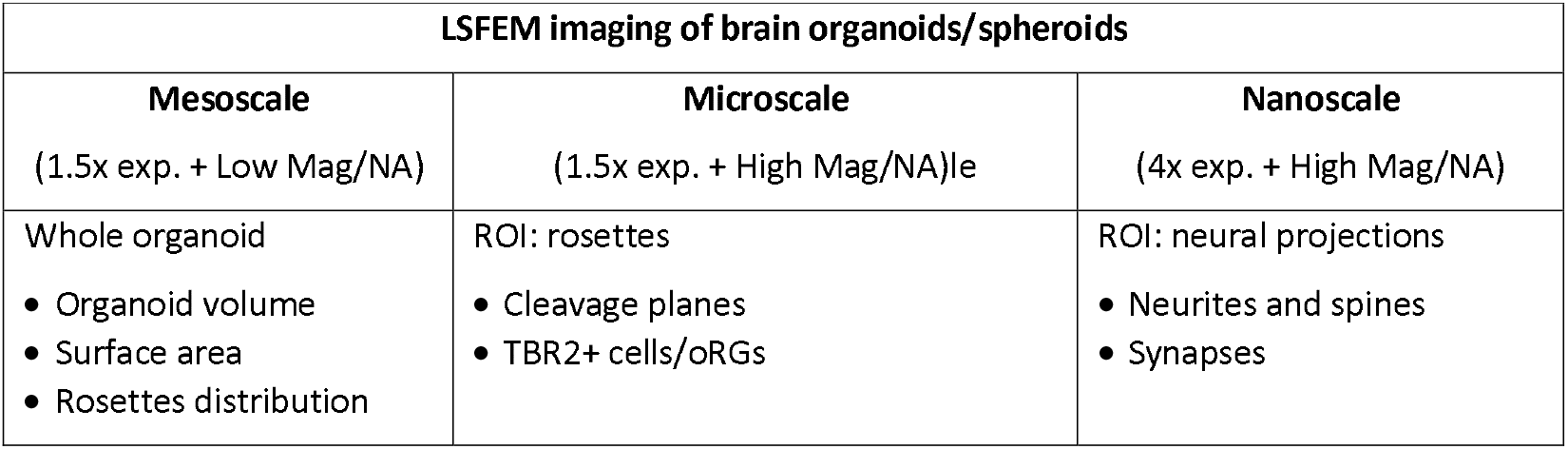
Imaging brain organoids at various scales with different combinations of the physical sample expansion (exp) and specific objectives lenses (Mag, magnification). An example of a region of interest (ROI) is given for each case, followed below by possible corresponding image analysis parameters. Imaging specific ROIs is especially relevant for the micro- and nanoscale due to the large amount of data that could be generated.

We employed mixed organoids containing 10% EGFP expressing cells (90% of iPSCs mixed with 10% of doxycycline-inducible EGFP-labeled iPSCs from the same genetic background prior to seeding them) and with all cell nuclei labeled by Hoechst. Use of our optimized sample preparation protocol (Bürgers et al., 2019; Stockhausen et al., 2020) allowed us to maintain the fluorescence of autofluorescent proteins, e.g. EGFP, by optimizing firstly the permeabilization of the sample and secondly digestion conditions and the expansion buffer, which avoided the necessity of antibody staining in this case. One month after the generation of the organoid the distribution of the EGFP cells was found to be not uniform throughout the volume as could be concluded from 2D images (Fig 3A-D). Rather, the labeled cells tended to form large clusters. Interestingly, such clusters of EGFP-labeled cells still existed after 14 months (Fig 3E-G).

**Figure 3.**
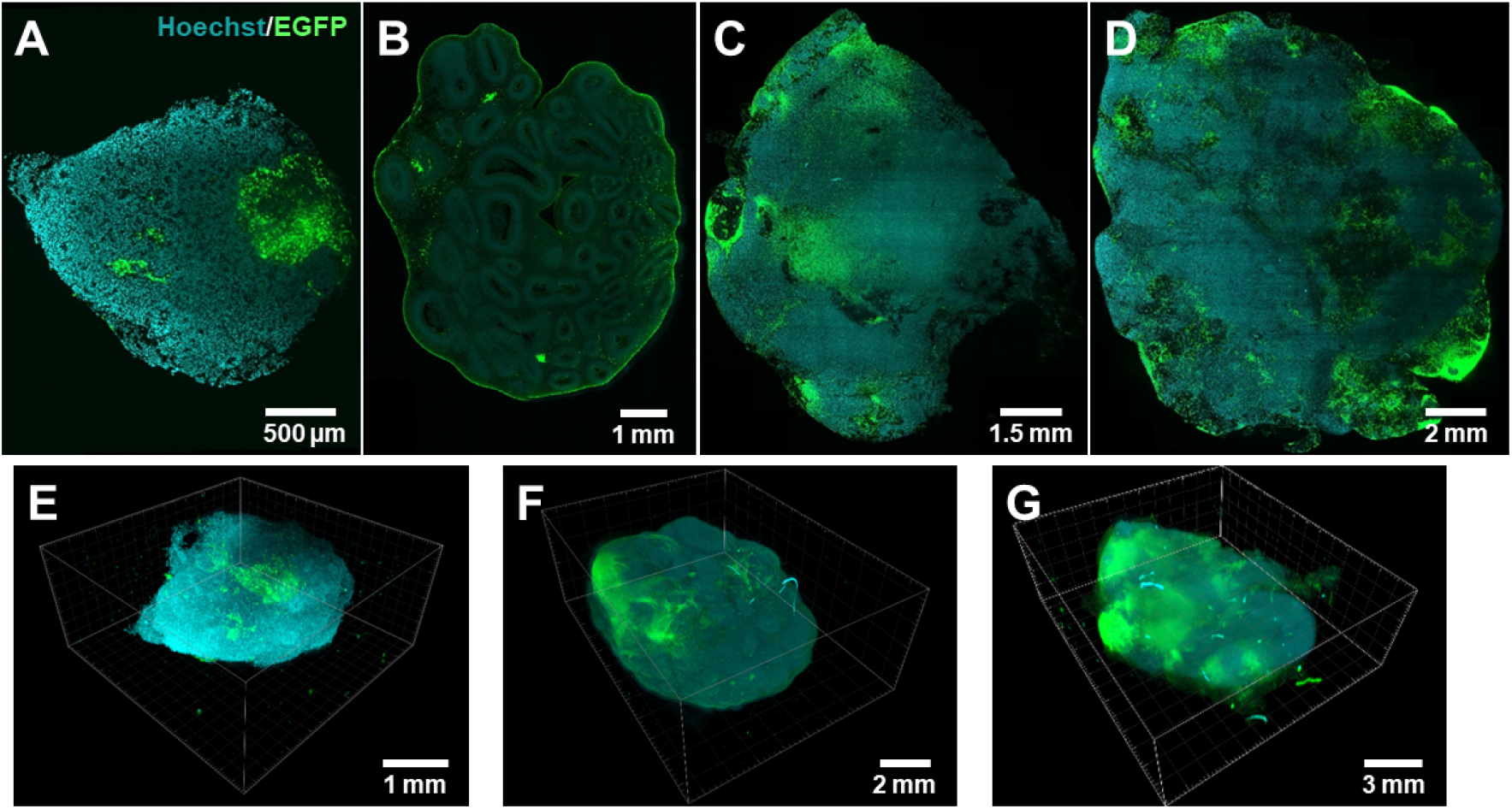
Development of chimeric brain organoids containing 10% EGFP expressing cells across a time span of 14 months. LSFM of cleared and 1.5-fold expanded brain organoids with cell nuclear staining containing 10% EGFP expressing cells (green) and nuclei are stained with Hoechst (cyan). (A) One month, (B) three months, (C) five months, and (D) fourteen months old brain organoids. The shown optical sections were taken at 646 μm, 1140 μm, 1830 μm, and 1119 μm depth, respectively. Image sizes were 3.6 × 3.6 mm^2^, 8.6 × 11.7 mm^2^, 13.1 × 14.9 mm^2^, and 16.6 × 17.3 mm^2^, respectively. (E) 3D view of the one month, (F) three months, and (G) five months old brain organoids, respectively. The imaged volumes corresponded to 3.6 × 3.6 × 1.6 mm^3^, 8.6 × 11.7 × 4.4 mm^3^, and 13.1 × 14.9 × 5.2 mm^3^, respectively.

##### LSFM and LSFEM allow meso-to nanoscale analysis in a single sample

The functional architecture of brain organoids extends over length scales from more than a centimeter to nanometers, and we became interested in devising an approach, which enables recording across these scales. Cleared and 1.5-fold expanded, complete brain organoids were imaged using a 10x objective lens (Fig. 3 and Figs. 4A, 4B). The use of 4-fold expansion, an 1.1 NA objective lens for imaging, an axial step size of 0.3 μm and subsequent deconvolution allowed visualization of selected sample regions at the 100 nm scale (Fig. 4C). Numerous cell somata and neurites with extensions up to hundreds of micrometers were visible and traceable. Close examination of magnified sample regions revealed numerous spine-like structures suggesting advanced differentiation and formation of neuronal connections (Figs. 4D, 4E and 4F).

**Figure 4.**
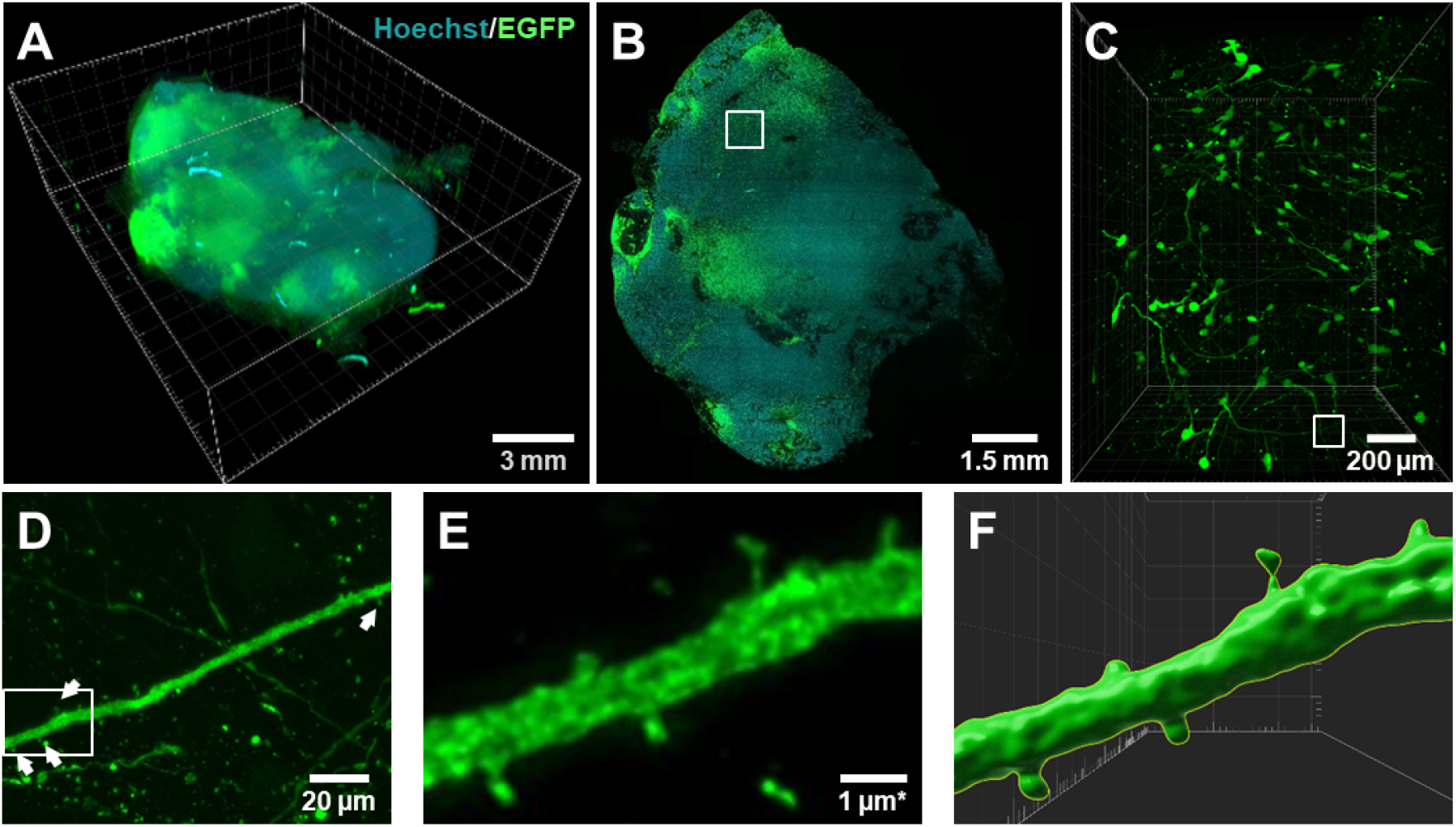
Five months old brain organoid containing GFP-positive cells imaged from the centimeter to the nanometer scale. (A) 3D view, volume 13.1 × 14.9 × 5.2 mm^3^. (B) Optical slice at a depth of 1.8 mm. The image was obtained using a 10x NA 0.3 objective lens. Size 13.1 × 14.9 mm^2^ (C) Rendering of a 3D stack with a volume of 1248 × 1548 × 1275 μm^3^ as marked in (B). The image was obtained using a 25x NA 1.1 objective lens in the same sample after 4-fold expansion. (D) Magnification of the region marked in (C), 185 × 132 μm^2^ revealing spine-like structures (see arrowheads). (E) Magnification of the region marked in (D). The adjusted scale bar considered the 4-fold expansion. (F) Surface rendering of the neural projection revealed spine-like structures. For (C) to (E) the shown image data were deconvolved. In total, 35 image stacks covering a total specimen region of 1248 × 1548 μm^2^ with a total depth of 1275 μm^3^, which was covered at an axial step size of 0.3 μm, were acquired from this organoid.

##### Qualitative and quantitative assessment of neuroepithelial architectures

Our approach enables detailed insights into the cellular architecture of brain organoids. Numerous structural parameters may be evaluated, when exploiting the fact that antibodies penetrate expanded tissue particularly well (Edwards et al., 2020). This also allows quantitative analysis, as we demonstrate here using neuroepithelial rosettes as example. These structures typically appear in cerebral spheroids, forming ventricular zone (VZ)-like areas, whose apical surface can be labeled with antibodies to the tight junction protein *zonula occludens protein 1* (Zo1). Zo1 immunofluorescence thus enables the evaluation of the topology of neuroepithelial rosettes and the ventricle-like space they could enclose (Fig. 5A and 5B). Fig. 5C shows the cropped apical surface across the whole organoid, demonstrating that these structures exhibit a large variation in size and shape. Some were closed structures, i.e. enclosing a ventricle-like lumen; others appeared relatively flat with a complex geometry and a sheet-like topology. Therefore, the measurement of the apical surface of the neuroepithelium appeared as an appropriate parameter for their characterization. The surface size distribution of the segmented structures of the organoid shown in Fig. 5A is displayed in Fig. 5C. Image data like that shown in Fig. 5C allowed the quantification of numerous parameters (Table 1), e.g. characterizing the apical surface topology (Fig. 5D). Mesoscale parameters were quantified imaging three different two months-old organoids.

**Figure 5.**
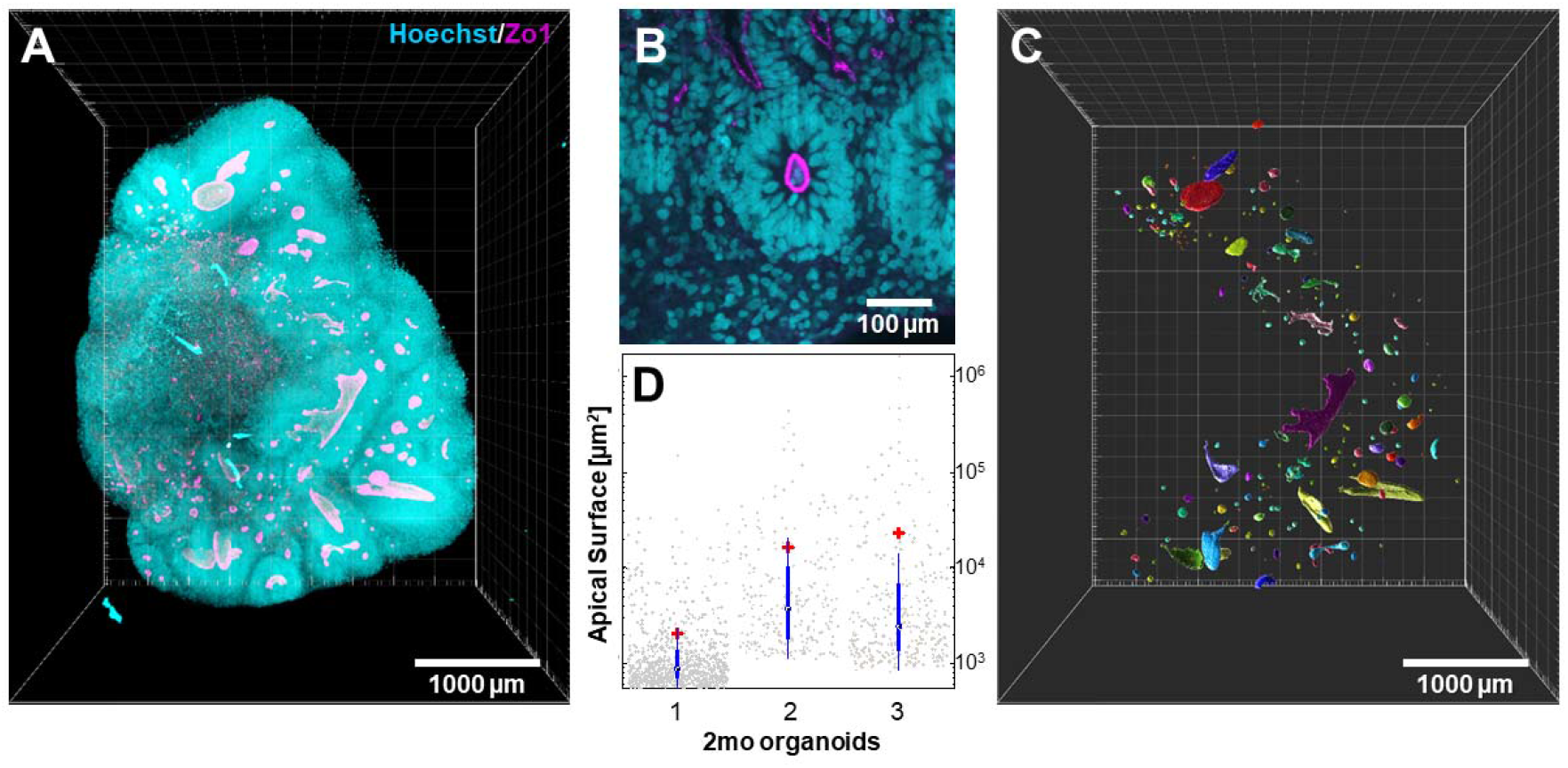
Labeling of the apical surface of neuroepithelial structures in a two months old brain organoid with an antibody against Zo1 (see Movie 1). Three different two months old organoids were imaged and evaluated to obtain the quantitative parameters as given below. (A) Cell nuclei (Hoechst, cyan) and Zo1 (magenta). The cyan fluorescent cell nuclei indicate the rough shape of the organoid. In this way, the total organoid volume (1.36 ± 0.55) × 10^10^μm^3^ and the total surface area of (1.2 ± 0.6) × 10^8^ μm^2^ was obtained. (B) Magnification of an optical section at a depth of 1 mm showing a closed apical surface revealing a VZ-like lumen inside a rosette. Considering that each rosette contains one apical surface, the average number of rosettes within the three whole organoids was evaluated yielding 485 ± 270 rosettes. (C) Cropped apical surfaces of the neuroepithelium inside the organoid shown in (A). Color labeling according to surface area, randomly generated. (D) Distribution of the apical surface areas of the three different two months old organoids appears as an appropriate parameter for their characterization, since not all structures enclose a volume. Mean values were indicated by the red cross. Overall mean is (1.66 ± 4.79) × 10^4^ μm^2^ and the median is 3710 μ m^2^.

##### Delineation of neural subpopulations

The combination of antibody staining and LSEFM allows straight-forward detection of neural subpopulations. As example we focused on outer radial glial (oRG) cells, a distinct population of neuronal progenitors in the developing human brain, which are located in the outer subventricular zone (oSVZ). These cells are essential for neurogenesis and expansion of the human cortex (Bershteyn et al., 2017). One option to identify oRGs is combined labeling with antibodies to the intermediate progenitor marker Tbr2 and the radial glia marker Sox2 (Fig 6). ORGs are characterized by expressing Sox2, but not Tbr2 in the outer region of VZs (see green arrows in Figs. 6A and 6C).

**Figure 6.**
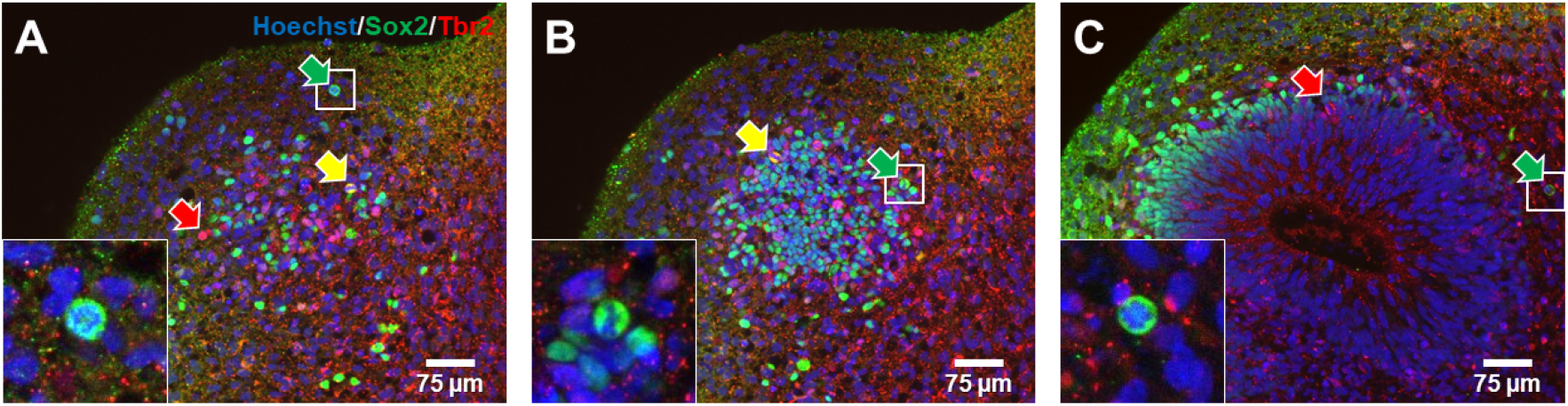
Identification of oRGs using the combination of Sox2 and Tbr2 labeling. Labeling of a rosette in a two months old brain organoid by Hoechst (blue), Sox2 (green) and Tbr2 (red). (A, B and C) show single optical slices taken at depths of 53 μm, 75 μm and 146 μm, respectively, in the sample. Cells being Tbr2 positive, but Sox2 negative (red arrows), both Tbr2 and Sox2 positive (yellow arrows) and Tbr2 negative, but Sox2 positive (green arrows) were marked. The latter ones are indicative for oRGs. Total area, 703 × 651 μm^2^.

However, only identifying Sox2 positive cells and their position is not sufficient for an unequivocal delineation and quantification of oRGs (Pollen et. al., 2019). Recent studies revealed that the vast majority of cells expressing HOPX also expressed the radial glia marker SOX2, but not the intermediate progenitor marker Tbr2 (Bhaduri et al., 2020, Pollen et al., 2019). Thus, HOPX is considered a useful marker for oRGs, and we applied it in three-month-old organoids. In Fig. 7, we show several examples for the identification of oRGs in different samples. Co-staining with an antibody to N-cadherin enabled the delineation of the apical surface of the neuroepithelium and thus a spatial relationship of the HOPX-positive cells to the cytoarchitecture (Figs. 7C and 7D).

**Figure 7.**
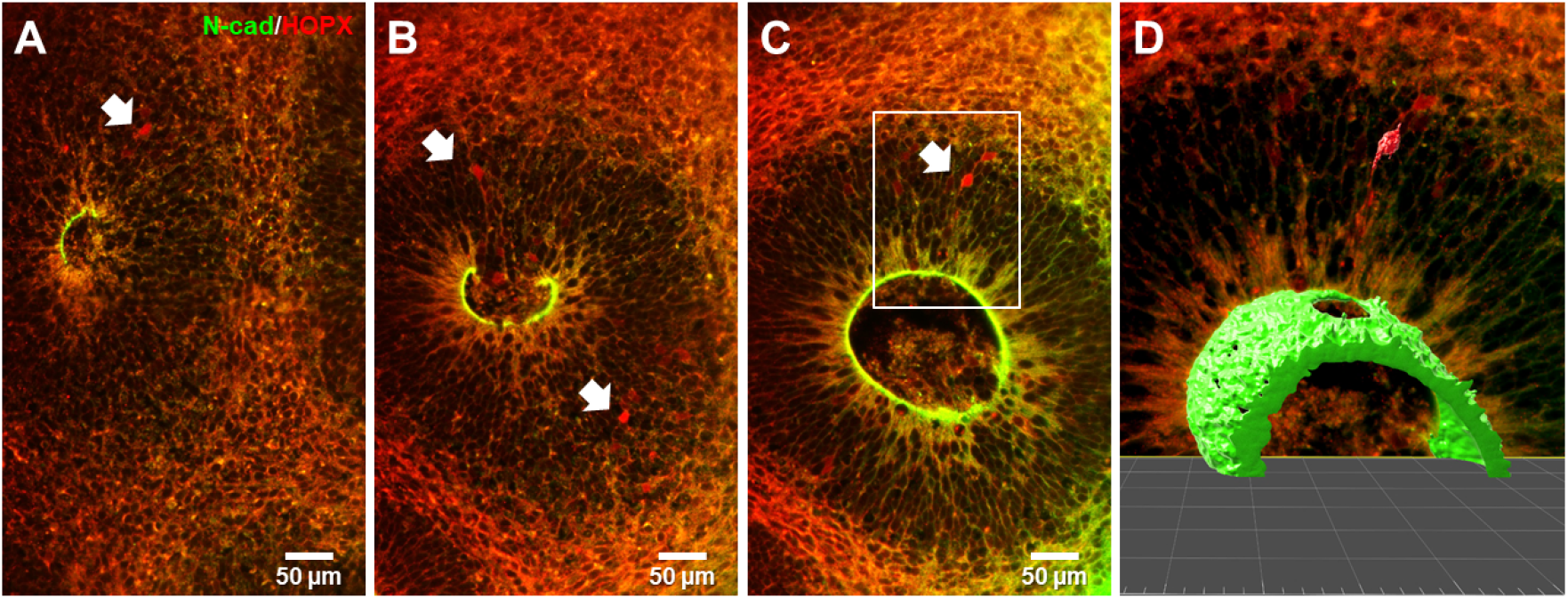
Definition of the 3D location of oRGs with regard to the VZ surface in three months old brain organoid labeled by N-cad (green) and HOPX (red). ORGs were marked by white arrows. (A) to (C), maximum intensity projection of 3 μm of optical sections at a depth of 200, 452, and 595 μm, respectively. (D) Surface rendering of oRG and VZ marked in (C).

#### Imaging of subcellular structures

##### Orientation of mitotic cleavage planes

A key parameter in neurogenesis during human brain development is the orientation of mitotic cleavage planes of neuronal progenitors with regard to the apical surface of the neuroepithelium (Fig. 8A). The orientation of the mitotic spindle modulates the orientation of the cleavage plane and, therefore, the position of the two daughter cells. The correct spindle orientation during the early stages of human corticogenesis is vital for accomplishing the right ratio between symmetric and asymmetric cell divisions. The prevalent number of actively diving neuronal progenitors during early cortical development exhibit a horizontal orientation (i.e., an angle of 0 to 30°) in relation to the ventricular surface cleavage plane, which leads to expansion of the cortical progenitor pool via symmetric cell division. Vertical (60 to 90°) and oblique (30 to 60°) mitotic cleavage planes start to become more predominant prior to neurogenesis (LaMonica et al., 2013; Yingling et al., 2008). This asymmetric mode of cell division results in the generation of two different daughter cells and leads to an increase in neuronal differentiation. Several studies have shown that a disrupted orientation of the mitotic cleavage planes led to abnormal corticogenesis reflected in various developmental phenotypes. Lancaster and colleagues performed one of the first landmark studies showing a crucial role of the shift in mitotic cleavage plane orientation and its effect on the development of microcephaly (Lancaster et al., 2013). In two more recent studies utilizing Miller-Dieker syndrome patient-derived organoids in comparison to controls, they have successfully shown that under disease conditions, there is a clear shift from the vertical to the horizontal plane of cell division of RGs without a significant increase in oblique planes, causing early neurogenesis and smaller size of patient-derived 3D cortical cultures (Bershteyn et al., 2017; Iefremova et al., 2017).

**Figure 8.**
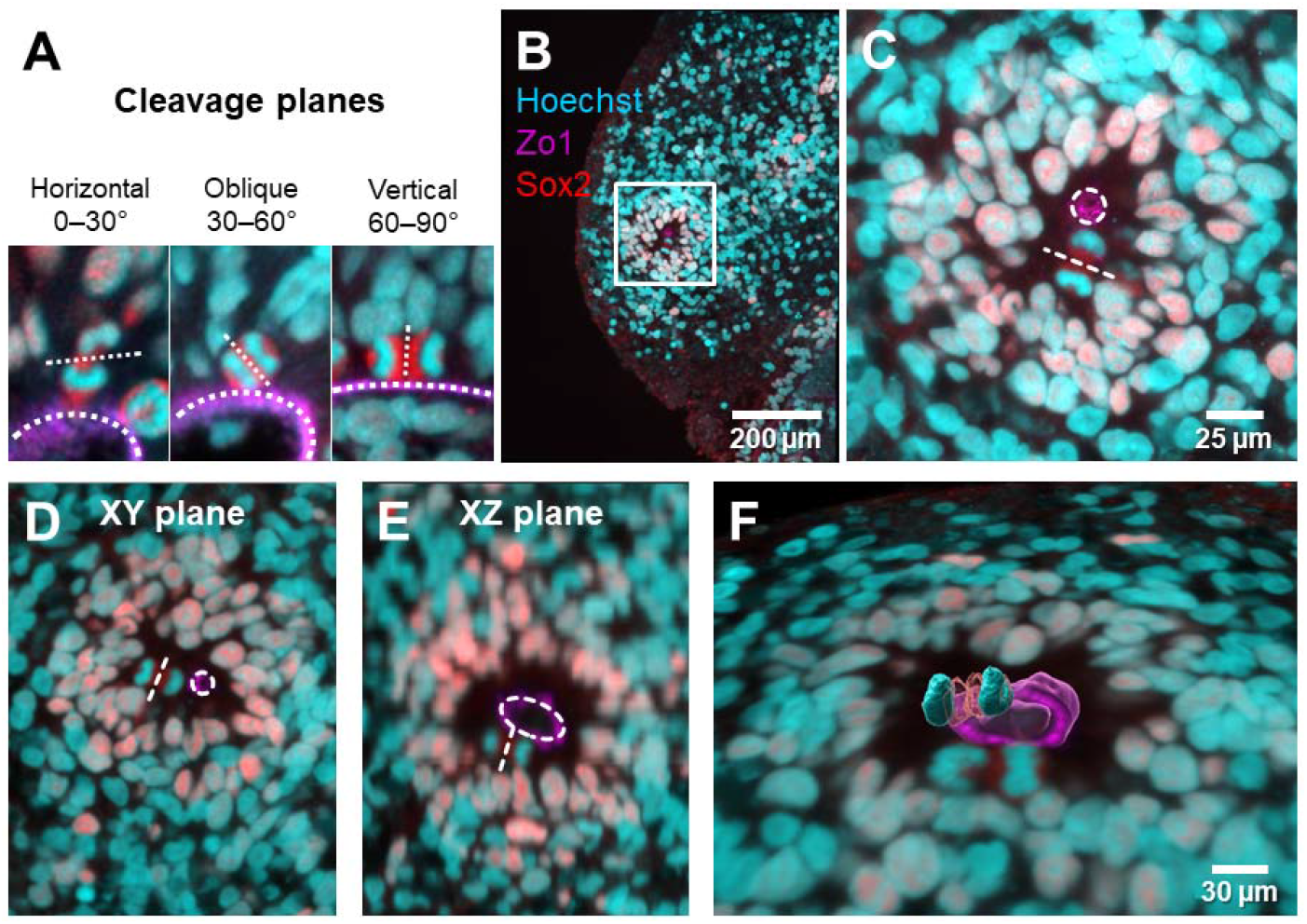
Analysis of cleavage planes. (A) Definition of cleavage planes with regard to the apical VZ lumen surface. (B) A two-month-old brain organoid was labeled by Zo1 (magenta), Sox2 (red) and Hoechst (cyan). Zo1 revealed the surface of a VZ lumen. Total area, 703 × 651 μm^2^. (C) Magnification of the region marked in (B). The orientation of the cleavage plane of a mitotic cell (dashed line) in relation to the lumen surface could be visualized. Total area, 277 × 277 μm^2^. (D, E) Using only 2D data may lead to a misinterpretation of the cleavage plane orientation. (F) The true cleavage plane orientation can only be deduced from 3D data (Movie 2).

We found the combination of DNA staining by Hoechst and immunofluorescence staining for Zo1 and the neural progenitor marker Sox2 suitable for delineating the orientation of cleavage planes with respect to the surface of the neuroepithelium (Figs. 8B and 8C). Importantly, our image analyses revealed that it is not always possible to assess the orientation of the cleavage plane with regard to the apical surface using 2D projections alone. As shown in Fig. 8D, the xy-section suggested a horizontal orientation of the cleavage plane of the mitotic cell with regard to the apical surface, whereas the xz-section of the very same cell nucleus indicated a vertical orientation. The evaluation of the true orientation requires the full 3D view (Fig. 8F).

##### Detection and spatial relationship of pre- and postsynaptic structures

The existence of functional neuronal connections in cerebral organoids has been reported previously (Paşca et al., 2015, Quadrato et al., 2017, Giandomenico et al., 2019). However, there are no reported studies using whole cleared organoids showing the co-existence of both pre- and postsynaptic proteins at synapses in a special manner (for review of current organoids imaging, see Brémond Martin et al., 2021). We used again chimeric organoids generated with 10% EGFP containing cells for such structures. Figure 9 shows a 14 months old organoid with a diameter of about 1.5 cm (Fig. 9A). Regions near the surface of the organoid contained numerous cells with neuronal morphology and long neurites (Fig. 9B). LSFEM with antibodies to the pre-synaptic protein synapsin1 (Syn1) and the post-synaptic protein Homer1 revealed diffraction limited spots within distances of 150 ± 61 nm (n=26), with a median value of 130 nm of each other along with neural projections (Figs. 9D and 9E).

**Figure 9.**
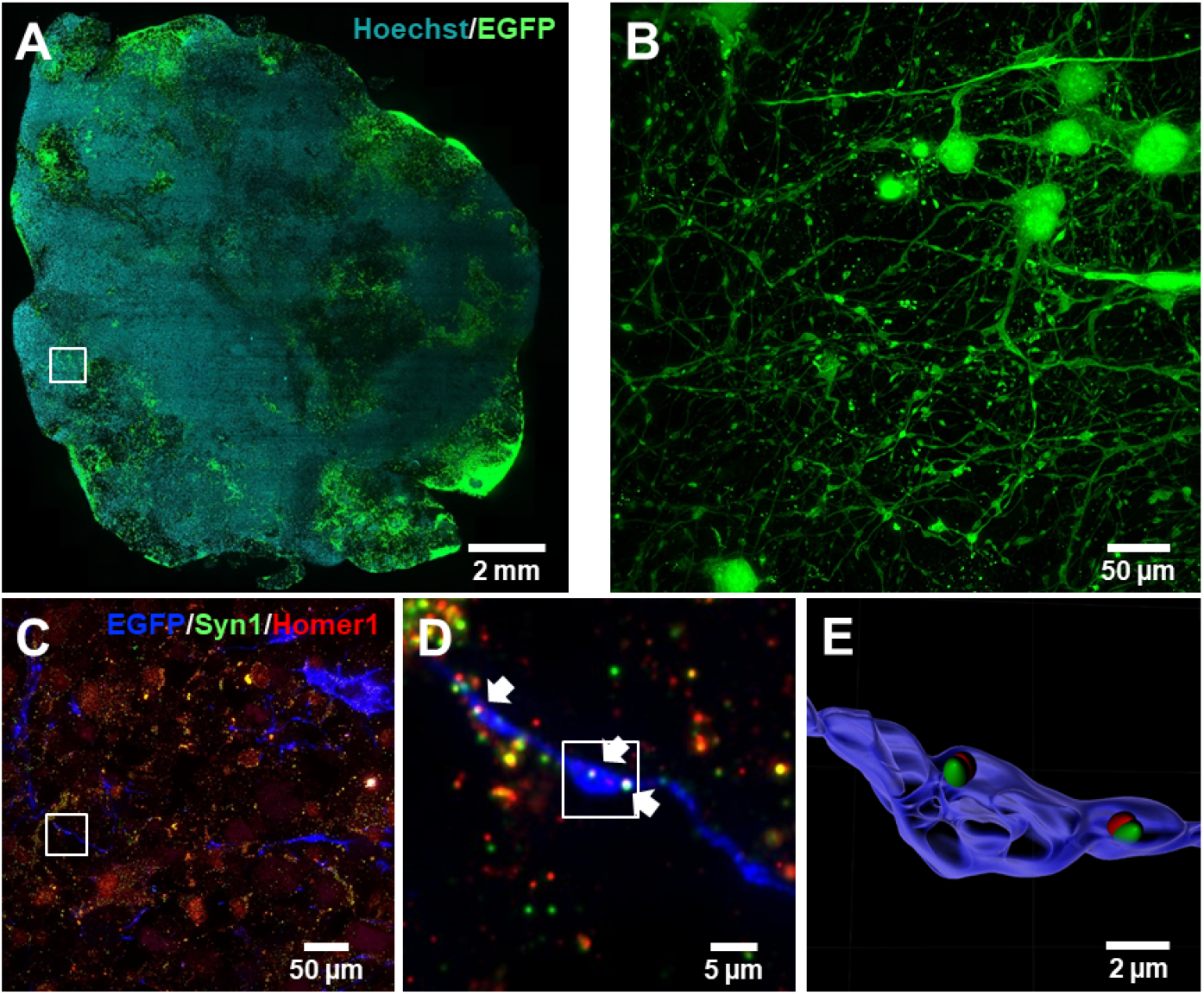
Pre- and postsynaptic structures in a 14 months old brain organoid. The organoid contained 10% EGFP expressing cells and was labeled by Hoechst and stained with antibodies to the pre- and post-synaptic proteins Synapsin 1 (green) and Homer 1 (red). For details see Movie 3. (A) Optical slice at a depth of 1.5 mm. This image was constructed using 285 single image tiles, which were acquired using a 10x NA 0.3 water immersion objective. Size 16.6 × 17.3 mm^2^ (B) MIP comprising 1000 images (about 300 μm in axial direction) acquired with an 25x NA 1.1 objective, size 547 × 547 μm^2^ after expanding the sample 4-fold, and after deconvolution. (C) MIP comprising 20 images (about 300 μm in axial direction). Here, EGFP is shown in blue and the pre- and postsynaptic proteins Synapsin 1 and Homer1 are shown in green and red, respectively. (D) Magnifications of the region indicated in (C), revealing the colocalization of pre- and post-synaptic proteins Syn1 and Homer1 along axonal boutons. Arrowheads indicates the synapses along neurites. (E) 3D reconstruction of the multisynaptic bouton showed in (D).

## Discussion

Cerebral organoids are opaque 3D structures with a size in the range of a few millimeters. Opacity and volume make light microscopic analysis difficult because the opacity impedes classical imaging with sufficient contrast and the organoid size prohibits the use of high-resolution optical microscopy. The latter requires objectives with a high NA, which generally have very short working distances. Classical imaging approaches using organoid slices miss the natural 3D features of such complex samples.

We demonstrate here the potential of LSFM combined with a clearing and expansion of organoids by a factor of 1.5 to 4-fold, which was recently already used for the examination of mouse brain sections (LSFEM, Bürgers et. al., 2019; Gao et. al., 2019; Stockhausen et. al., 2020). This new approach was refined by a careful preparation of the organoid tissue exploiting permeabilization using CHAPS instead of Triton-X (Zhao et al., 2020). These adaptations enabled especially the preservation of the activity of autofluorescent proteins and staining of complete, large organoids using commercial antibodies with high efficiency. We demonstrate that this approach allows the detailed analysis of organoids with a size up to 15 mm.

The preservation of the fluorescence of autofluorescent proteins during the expansion procedure enabled us to follow the location and fate of selected cell types during the course of development over a time period of up to 14 months. The use of a fraction of cells expressing fluorescent proteins when generating the organoids in addition to nuclear staining allowed a deeper analysis of the location and distribution of cells groups within the volume. Thereby, we confirmed the well-known observation (Quadrato et al., 2016; Quadrato et al., 2017; Qian et al., 2019; Velasco et al., 2019) that the development of organoids varied largely due to the batch-to-batch variability.

The critical details of neuronal connectivity occur on length scales of about 100 nm. Such small structures can optically only be resolved using super-resolution light microscopy. We have already demonstrated that LSFEM can yield effective super resolution laterally down to less than 100 nm and axially down to 300 nm. Thereby, individual synaptic connections can be identified (Bürgers et al., 2019). Achieving this requires clearing followed by an expansion of the sample by a factor of 4 and subsequent high-resolution imaging. Further improvement of image resolution may be achieved by deconvolution techniques. This approach allowed us here to detect single spine like structures in a five-month-old brain organoid. Usually, imaging at such a high resolution is feasible but not applicable for large sections of organoids due to the immense amount of data produced. An organoid of 1 mm^3^ original size would yield after expansion an object of 64 mm^3^ size. Imaging that structure at a resolution of 100 nm laterally and 300 nm axially at 16 bit considering the Nyquist theorem would yield a data set of 340 TB. Current data processing workstations are at or beyond their computational limit when handling such amounts of data. Therefore, imaging at a mesoscopic scale is used to locate specific regions, for which super-resolution data shall be obtained. Notably, this allows examining a single specimen on length scales from centimeter to 100 nm corresponding to 5 orders of magnitude.

A hallmark of developing cerebral organoids is the generation of rosette-forming neuroepithelial structures with an apical-basal polarity. The space enclosed by these structures has been shown to form a ventricle-like system, which can span across large volumes of the organoid (Di Lullo and Kriegstein, 2017). As these areas correspond to a pendant of the neurogenic ventricular zone in vivo, their qualitative and quantitative assessment is of great importance. The study of these structures was straightforward using LSFEM because they are immediately visible in the 3D data we acquired. The lumina of VZs could well be visualized and analyzed using additional staining against Zo1 or N-cadherin.

A further determinant of brain organoid structure are oRGs. These cells can be stained using antibodies against Tbr2, Sox2, and HOPX (Pollen et al., 2019). The identification of oRGs is generally challenging because of the low abundance of this cell type in brain organoids. Therefore, it is even more problematic, if only 2D sections are employed, because the mapped volume is quite small. The use of the complete 3D image stack clearly improved the chance to detect this important cell type. For the success of these experiments and to achieve a labeling with high contrast it was especially important to employ a new permeabilization strategy before labeling, namely to use CHAPS instead of Triton X (Zhao et al., 2020). This approach yielded a much better penetration of antibodies to their target sites inside the mature, rather large organoids.

The orientation of cleavage planes of dividing cells with regard to the VZ surface is important for the growth properties of a brain organoid. We noted that only evaluating the 3D data revealed the correct orientation of cleavage planes with regard to the lumen surface. Availability of 3D data avoided possible misinterpretations compared to using only 2D optical sections. Furthermore, we noticed that the use of Sox2 is sufficient to visualize the orientation of cleavage planes with our method, without the need for specialized antibodies like pVimentin.

In contrast to electron microscopy, LSFEM is compatible with multicolor fluorescence imaging, thus enabling molecular contrast for diverse neuronal populations and nanoscale resolution within a single, large tissue preparation. Here, we performed triple color staining of the brain organoid using EGFP and pre-and postsynaptic markers in order to identify synapses unambiguously. Using LSFEM, we succeeded here for the first time to detect both pre-synaptic Synapsin1 and post-synaptic Homer1 within distances of 150 nm of each other along with neural projections clearly proving the existence of synapses in a fourteen-month-old organoid. In principle, our technique provides a sufficient optical resolution to allow the identification of different kinds of spines, e.g. mushroom spines or stubby spines, in 3D. In the future for such experiments it would be helpful to label dendritic structures using MAP2.

In summary, we demonstrated that combination of LSFM and ExM - LFSEM - allows to image mature brain organoids *in toto* and down to synaptic resolution, when they are combined with careful specimen preparation preserving autofluorescent proteins and finally optimizing imaging results using deconvolution. Thus, LSFEM is optimally suited for the analysis of brain organoid development.

## Materials and Methods

### Generation of iPSC-derived 3D organoids

Organoids were generated along previously established protocol with slight modifications (Iefremova et al., 2017). In brief: on day 0 of organoid culture, iPSCs were dissociated into single-cell suspension using TrypLE Express, followed by plating 18000 cells in each well of an ultra-low-attachment round-bottom 96-well plate in StemFlex medium with 50μM of ROCK inhibitor Y-27632. In order to visualize the EGFP-positive cells, iPSCs from the same genetic background with and without Dox-inducible EGFP construct were mixed in the ratio 10/90, respectively. Organoids were fed every other day for up to 5 days with StemFlex media and then transferred to low-adhesion 6-cm plates in the neural induction medium containing 50% Neurobasal, 50% DMEM/F12 supplemented with 1x N-2 supplement, B-27 supplement, and glucose (0.4mg/mL). Neural induction medium was supplemented with 1% NEEAE, 1% GlutaMax TM, LDN-193189 (180nM), A8301 (500nM) and XAV (10 μg/ml) before medium change. Next, in 5-6 days, the medium was changed to the neural differentiation medium containing 50% Neurobasal, 50% DMEM/F12 supplemented with 1x N-2 supplement, B-27 supplement, glucose (0.4mg/mL), cAMP (0.15 μg/mL), 1 % NEEA, 1 % GlutaMax TM. Organoids were embedded within the next 5 days in Matrigel and further cultured on a cell culture shaker with a medium change every 2-4 days until the day the cultures were fixed for further analysis. From D35 onwards the medium was changed to 50% Neurobasal, 50% DMEM/F12 supplemented with 1x N-2 supplement, 1x B-27 supplement, and glucose (0.4 mg/mL) cAMP (0.15 μg/mL), Insulin, 1% Matrigel and 20 ng/ mL BDNF and 10 ng/mL GDNF.

### Generation of iPSC-derived 3D cortical spheroids

The generation of cortical spheroids was performed by using a modified protocol from (Paşca et al. 2015). In brief, iPSCs were dissociated into single cell-suspension with StemPro Accutase (Thermo Fisher Scientific) and spheroid formation was performed by transferring 1.5 × 106 iPSCs (5,000 cells/microwell) into AggreWell 800 plates (Stem Cell Technologies) in spheroid medium (50% DMEM-F12 Glutamax, 50% Neurobasal, 1:100 B27, 1:200 N2, 1:200 MEM-NEAA, 1 mM L-Glutamine, 10 μg/ml Insulin and 1:1000 b-mercaptoethanol) supplemented with the two SMAD pathway inhibitors, dorsomorphin (1 μM, Sigma Aldrich) and SB-431542 (10 μM, AxonMedChem), as well as with the ROCK inhibitor Y-27632 (10 μM, Hiss Diagnostics). As described for organoid generation, iPSCs from the same genetic background with and without Dox-inducible eGFP construct were mixed in the ratio 10/90. For the first five days, the spheroid medium without ROCK inhibitor was changed daily. Afterwards, the spheroids were transferred into a CERO tube and cultivated in a rotating CERO table-top bioreactor (CERO 3D bioreactor, OLS). From day 5 to day 12, spheroids were fed every other day. On day 12, spheroid medium containing bFGF (10 ng/ml, Biotechne) instead of SMAD inhibitors was used for four days. From day 16 on, spheroids were maintained in unsupplemented spheroid medium with medium changes every other day.

### Generation of the mixed 3D cultures containing doxycycline-inducible EGFP-labeled iPSCs

In order to generate mixed 3D cortical cultures (both organoids and spheroids), 90% of iPSCs were mixed with 10% of doxycycline-inducible EGFP-labeled iPSCs from the same genetic background prior to seeding them. The establishment of the EGFP line was performed with the use of transcription activator-like effector nucleases-mediated targeting to the AAVS1 site as previously described by (Qian et al., 2014). After plating the mixed cultures on the respective plates, according to the protocols listed above, 3D cultures were maintained under the same conditions as iPSC-derived cultures containing 100% of unlabeled cells.

### Mesoscopic Imaging

Mesoscopic imaging and analysis yield information on the topology of complete organoids, which is helpful, for example, to analyze batch-to-batch differences. In order to allow rapid imaging at a sufficient resolution, samples were first fixed and permeabilized. Subsequently, staining of specific cell types was performed using nuclear staining and commercially available antibodies, e.g. against the tight junction marker Zo1. Now the ExM protocol started with linking the basic amine groups to a polyacrylamide gel that is formed within the sample. Finally, the tissue was digested (or “homogenized”) using proteinase K (Prot-K), which rendered the sample transparent. Placing the sample into phosphate buffered saline (PBS) leads to an expansion by a factor of 1.5.

For imaging, the digested specimen was fixed on a coverslip with poly-L-lysine to avoid movements during the measurement. Then, the coverslip was inserted into the sample holder and placed into the sample chamber filled with PBS solution. Before image acquisition, a visual inspection of the sample was performed to verify a successful sample preparation. Then, the samples were analyzed using LSFM employing an 10x objective lens with a NA of 0.3 and an effective field of view of (998 μm)^2^. The achieved real and effective optical resolutions were given in Table 2.

**Table 2.** True and effective optical resolutions of imaging at various scales

| Scale | Objective lens | True optical resolution |  | Effective optical resolution |  | Data size per 1 mm<sup>3</sup> sample |
| --- | --- | --- | --- | --- | --- | --- |
|  |  | Lateral | axial | Lateral | axial |  |
| Mesoscale | 10x, NA 0.3, WI | 1.2 $\mu\text{m}$ | 17.8 $\mu\text{m}$ | 0.8 $\mu\text{m}$ | 11.8 $\mu\text{m}$ | 5.5 GB |
| Microscale | 25x, NA 1.1 WI | 300 nm | 1100 nm | 200 nm | 800 nm | 500 GB |
| Nanoscale | 25x, NA 1.1 WI | 300 nm | 1100 nm | 100 nm | 300 nm | 8 TB |

Due to the large size of the sample, imaging in a mosaic fashion was needed in order to image the whole organoid. Finally, the data were stitched using the stitching plugin of FIJI, and imported into Imaris (Bitplane AG, Zurich, Switzerland) for further analysis.

### Microscopic Imaging

For imaging at the microscopic scale the organoids were prepared as described above, yielding a transparent, 1.5-fold expanded specimen. Now, however, they were examined with a high-resolution, long-distance objective with an NA of 1.1 enabling an optical resolution of about 0.3 μm laterally and 1.1 μm axially. This resulted in effective resolutions of 0.2 and 0.7 μm, respectively, when considering the sample expansion rate (Table 2). Thus, the lateral resolution increased by a factor of ≈4 and the axial resolution by a factor of ≈15 compared to mesoscopic imaging. This made a structural characterization at cellular length scales possible. Imaging of complete organoids at this resolution would produce about 500 Gigabyte data per 1 mm^3^ and per channel, which would require high end image processing workstations for analysis. Therefore, imaging of complete organoids at this resolution is often not advisable, although it is principally possible. Rather, certain ROIs should be selected in the mesoscale data for subsequent analysis at the microscopic scale.

### Nanoscopic Imaging

For imaging at the nanoscopic scale, organoids were prepared as described above yielding transparent, 1.5-fold expanded specimen. Then, the buffer solution, into which the sample was placed, is replaced by bi-distilled water. This leads to an approximately four-fold expansion compared to the original sample size. Such samples were examined with a high-resolution, long-distance objective (NA 1.1) enabling an optical resolution of about 0.3 μm laterally and 1.1 μm axially. This yielded effective super-resolution of 0.1 and 0.3 μm, respectively, when considering the sample expansion (Table 2), which made a structural characterization at subcellular length scales possible. The resolution can further be improved by 3D image deconvolution. We exemplified this imaging approach, first, to study spine formation in brain organoids. To this end, we employed five-months and fourteen-months old brain organoids containing sparse cells expressing EGFP. The application of 3D image deconvolution improved the level of sample detail even further. We acquired 3D image stacks which was covered at an axial step size of 0.3 μm. The axial step size was adjusted such that deconvolution using Huygens (Professional version 17.04, Scientific Volume Imaging, The Netherlands) could optimally be performed.

### I mm unochemistry

Immunohistochemistry was optimized from standard protocols. In brief, the fixed 3D cultures were first permeabilized using CHAPS in the permeabilization buffer (1xPBS, 0.5% CHAPS) on a shaker at 37°C. The time varies depending the size of the sample, e.g. 1 hr. for a one-month-old sample. After permeabilization, samples were washed three times with 1xPBS at room temperature (RT). To prevent from unspecific binding of the primary antibody, the samples were incubated with blocking buffer (1x PBS, 5% normal goat serum, 0.3% TritonX-100, 0.02% sodium azide) on a shaker ON at room temperature. After blocking, the sections were incubated for ON in primary antibody (SOX2, R&D Systems, MAB2018; TBR2, Abcam, ab23345; HOPX, Santa Cruz, sc398703; HOPX, Proteintech, 11419-1-AP; N-cadherin, BD Biosciences, 610921; Synapsin1, Synaptic Systems, 106 103; Homer1, Synaptic Systems, 160 004) on a shaker at +4°C. The following day, slices were washed at room temperature in blocking buffer three times for 30 min and incubated ON in secondary antibody (Alexa550 anti-ms, 43394, sigma-Aldrich; Alexa555 anti-ms, A21424, Thermo Fisher Scientific; Alexa555 anti-rb, A21429, Thermo Fisher Scientific; Alexa594 anti-gp, A11076, Thermo Fisher Scientific; Atto647N anti-ms, 50185, Sigma-Aldrich; Atto647N anti-rb, 40839, Sigma-Aldrich) on shaker at +4°C. For nuclear staining, all samples were stained using Hoechst 33342 (H3570, Invitrogen).

### Sample preparation

The expansion microscopy protocol was adopted from (Chozinski *et al*., 2016). The immunostained organoids were incubated with 2 mM methylacrylic acid-NHS (Sigma Aldrich) linker for 24 hours on a shaker at RT. After washing three times in PBS, the organoids were incubated 16 hours in the monomer solution (8.6% sodium acrylate, 2.5% acrylamide, 0.15% N,N’-methylenebisacrylamide, and 11.7% NaCl in 1× PBS) on a shaker at +4°C.

The gelling solution was prepared by adding 4-hydroxy-TEMPO (0.01%), TEMED (0.2%) and ammonium persulfate (0.2%) to fresh monomer solution. During gelling, the organoids were placed in a 24-well plate on ice to avoid early polymerization. After applying the gelling solution, samples were put on a shaker at +4°C for 5 minutes and then be transferred to the gelling chamber, followed by 3 hours incubation at 37°C. After the gel formation, the samples were incubated at 37°C in the digestion buffer (50 mM Tris, 1 mM EDTA, 0.5% Triton-X100, 0.8M guanidine HCl, and 16 U⍰ml of proteinase K; pH 8.0), exchanging the buffer every 24 hours. In general, a two m.o. organoid takes about two complete days to be completely digested. After digestion, the buffer was removed and the samples were washed three times with PBS.

For microscopic examination, the expanded gel sample was fixed on the bottom coverslip of the imaging chamber with poly-L-lysine to avoid movements during the measurement, and the chamber was filled with PBS.

### Data processing

3-D stacks of raw 16-bit images were processed using custom-written MATLAB scripts, which allowed parallel data processing (Gonzalez et al., 2009). In a first step, the intensity histograms were adjusted to homogenize brightness and contrast throughout the complete data set.

Complete 3D representations of the samples were possible after several 3D data sets we stitched together using FIJI (Schindelin et al., 2012), and the stitching plugin of (Preibisch et. al, 2009). In order to optimize the stitching process, especially when datasets exceed the available RAM memory of the workstation, the process was performed in two steps. First, substacks of the 3-D data sets were created using a FIJI script. Each substack contained about 15% of the information located in the center of the full stack. Secondly, each substack was stitched to its respective neighboring substack yielding the best overlap in terms of the cross-correlation measure. Based on the localization information of each substack after stitching, the full 3D stacks were stitched.

A final step to improve the contrast throughout the 3-D data was performed after stitching. This was done to compensate for possible intensity variations of the sample in the axial direction. To this end, a histogram equalization was performed in every image plane of the stitched data set. For calculation of z-projections, the maximum intensity projection algorithm of Fiji was used.

### Deconvolution

As outlined in the results section, selected image stacks were spatially deconvolved using Huygens (Professional version 21.10, Scientific Volume Imaging, The Netherlands). Deconvolution was performed using theoretical PSFs, based on microscopic parameters, or a measured PSF determined by analysis of fluorescent microbeads embedded in 1% agarose gel. The classical maximum likelihood estimation algorithm was used, and a SNR value between 12 and 20 for a maximum iterations between 60 and 100 were selected.

The 3-D representation of the data was achieved using the Surpass view in Imaris (Version 9.7.2 Bitplane Inc., Zurich, Switzerland). Data processing was performed on a workstation equipped with two Intel Xeon Platinum 8160 CPU (2.1 GHz, 24 cores), 512 GB memory, and an Nvidia Quadro P5000 GPU (16 GB GDDR5X) running under Windows 10 Pro.

## Acknowledgements

We gratefully acknowledge expert support by the micromechanical workshop of the Institute of Physical and Theoretical Chemistry of the University of Bonn under the guidance of Daniel Poetes.

## Data Availability

The datasets generated during the current study comprise numerous Terabytes of multichannel images and were processed and evaluated using Imaris (Bitplane). Access can be obtained upon reasonable request from the corresponding author.

